# White Matter Connectometry Among Individuals with Self-Reported Family History of Drug and Alcohol Use Disorders

**DOI:** 10.1101/644054

**Authors:** Abigail B. Waters, Kayle S. Sawyer, David A. Gansler

## Abstract

Heredity is an important risk factor for alcoholism. Several studies have been conducted on small groups of alcohol naïve adolescents which show lowered fractional anisotropy of frontal white matter in FH+ groups. We sought to compare large FH+ and FH-groups using white matter connectometry, as opposed to the previously used global tractography method, as it is more sensitive to regional variability. Imaging and behavioral data from the Human Connectome Project (WU-MINN HCP 1200) was used. Groups of participants were positive (n=109) and negative (n=109) for self-reported drug and alcohol use disorders in at least one parent. Groups were matched on gender, age, education, current alcohol usage, and alcohol use disorders (AUD). Connectometry was performed on diffusion MRI in DSI-Studio using q-space diffeomorphic reconstruction, and multiple regression was completed with 5000 permutations. Analyses showed decreased major tract (>40 mm) connectivity in the FH+ group in left inferior longitudinal fasciculus, bilateral cortico-striatal pathway, left cortico-thalamic pathway, and corpus callosum, compared to the FH- group. For cognitive tasks related to reward processing, inhibition, and monitoring, there were a number of interactions, such that the relationship between identified networks and behavior differed significantly between groups. Positive self-report of family history of alcoholism was associated with decreased connectivity in reward signaling pathways, controlling for alcohol consumption and AUD. This is the first connectometry study of FH+, and extends the neural basis of the hereditary diathesis of alcoholism beyond that demonstrated with global tractography. Regions associated with FH+ are similar to those associated with AUD.

## Introduction

Family history of drug and alcohol abuse is highly associated with more severe, more recurrent alcohol dependence subtypes (Moss *et al.* 2007) and substance use disorders (Merikangas & McClair 2012), which remain a critical global health concern and disease burden (Whiteford *et al.* 2013). The impact of family history is further supported by extensive research in twin studies, demonstrating a genetic conferral of vulnerability when first-degree relatives have history of drug and alcohol problems (Walters 2002; Agrawal & Lynskey 2008). What remains a topic of continued exploration are the *specific mechanisms* of biological vulnerability in this population and how they relate to clinically relevant behaviors that interact with environmental stressors. Identifying brain structures underlying functional systems that differ among individuals with family history may help to explain how vulnerabilities are conferred, and identify critical targets and periods of intervention.

Identifying aspects of brain white matter structural connectivity vulnerabilities may be especially helpful, given that the developmental trajectory of these structures see the largest periods of growth when risk for alcohol and drug first use is highest (Bava & Tapert 2010). Longitudinal comparisons between young children with a family history of substance use disorders and healthy controls (Corral *et al.* 2003) show persistent neuropsychological deficits on a task of set-shifting in the presence of reinforcement (e.g., Wisconsin Card Sort Task) at three-year follow up. These findings underscore the clinical importance of examining biological vulnerabilities in reward processing pathways, given the longstanding effects on cognition. A study of alcohol-naive adolescents (Herting *et al.* 2011) has identified decreased integrity of white matter microstructure in the anterior limb of the internal capsule and superior longitudinal fasciculus associated with family history of drug and alcohol problems. These structural differences were significantly associated with reduced contra-lateral, fronto-cerebellar connectivity during reward-based decision making task performance, executive monitoring, and inhibition. A second study in a similar population found that reduced integrity of white matter in the inferior longitudinal fasciculus and optic radiations mediated group differences between those with and without family history on a task of reward processing (Herting *et al.* 2010). There have been consistent findings for decreased fronto-cortical and striatal white matter (i.e., corona radiata, corpus callosum, thalamic radiation, occipitofrontal fasciculus) when positive family history is defined as any substance use behaviors (Acheson *et al.* 2014) among adolescents and young adults. In a single study (Squeglia *et al.* 2014), high functioning alcohol-naïve early adolescents with positive family history showed increased white matter integrity across interhemispheric, association, and fronto-subcortical projection fibers. Although these findings may seem contradictory, in early adolescence accelerated maturation of these pathways has been associated with risk taking behaviors (Berns *et al.* 2009), and may indicate general dysregulation of growth trajectories within this population. While there have been discrepancies among studies when identifying specific tracts, individuals with positive family history consistently show effects on white matter integrity in reward processing circuits.

Taken together, this suggests that the biological differences between those with and without a family history of chemical dependency (i.e., drug and alcohol addiction) precede both onset of substance use and the maturation of white matter. This may reflect different developmental trajectories for those with and without family history of drug and alcohol disorders, and what remains unclear is whether these differences reflect white matter development that is slowed or stunted; essentially, do these vulnerabilities persist into adulthood? Therefore, examining white matter at a developmental stage where it is expected to be more mature may better characterize which differences are most impactful. Acheson and colleagues (2014) compared the white matter microstructure in healthy controls and individuals with positive family history (FH) across two age groups: youth (ages 10-14; *N* = 80) and young adults (ages 18-30; *N* = 25). While there were fewer significant findings among adults, suggesting a slowed trajectory, differences in diffusion protocols, demographic variables, and sample size prevented the direct comparison of groups or testing of age effects. Further exploration regarding the impact of family history on white matter in young adulthood is warranted in light of these limitations.

Traditional tractography in white matter studies utilize fiber tracking algorithms and “end-to-end” measurements of white matter integrity (Hagmann *et al.* 2010). This means that anisotropy measures (e.g., fractional anisotropy) are generated through the length of a tract, and only using fibers that connect to both identified ends. Typically, these measures are considered a reflection of the integrity of white matter tracts, and communicate the degree to which water is differentially restricted within a *single* direction. However, fractional anisotropy cannot differentiate between axonal architecture and myelin, especially when fibers cross within the same voxel (Soares *et al.* 2013). In contrast, a local connectome analysis measures the connectivity between *adjacent* voxels within white matter bundles, defined by the density of the diffusing spins, and shows changes in tract compactness or axonal density (Yeh *et al.* 2016). To generate the local connectomes used in group connectometry, image reconstruction maintains the spin distribution function (Yeh *et al.* 2010, 2011) which has a number of advantages.

Findings in adolescent samples have additionally been limited by the high presence of crossing fibers in these cortico-striatal regions, which may mask specific subcortical white matter structures when examining the average integrity along an entire tract. *Group connectometry* analyses can characterize variability within white matter tracts in the context of a local connectome, and identify clinically meaningful differences. Because it is not restricted by predetermined end points, this approach reduces noise from unrelated white matter branches. inflammation, and edema (Zhang *et al.* 2013). Additionally, the maintenance of spin directions in reconstruction mean that SDF based measures are better able to distinguish crossing fibers, and less impacted by partial volume effects, which are particularly salient when examining potential differences in cortico-striatal pathways implicated in reward processing (Rolls 2000), and dense connections between cortex and the corpus callosum. *In vivo* comparisons between traditional diffusion measurements (i.e., fractional anisotropy) and SDF based measures (i.e., quantitative anisotropy), show a reduction in false tract identification of approximately 50% (Yeh *et al.* 2014). Its increased sensitivity, compared to region-of-interest approaches, mean that it is particularly well suited to identify underlying vulnerabilities in populations where there may not yet be observable behavioral differences.

To our knowledge, this is the first group connectometry study investigating the effect of family history on SDF based measurements of white matter. We have chosen this approach because we believe that the density based measurements described above will be better able to characterize vulnerabilities conferred by parental drug and alcohol abuse disorders. By examining this relationship in adults, we hope to better characterize vulnerability during more advanced white matter development. Additionally, we will investigate the relevance of our findings as they relate to cognition in order to characterize the clinical impact of these findings. To answer these questions, we present the following hypotheses:

### Hypothesis 1

Compared to the individuals without family history (FH-), the individuals with family history (FH+) will have decreased connectivity along ventral tractography implicated in reward or inhibitory processes (i.e., cortico-striatal and fronto-cerebellar), as measured by measures of spin distribution function.

### Hypothesis 2

The relationship between measures of tract connectivity and clinically relevant behaviors (i.e., reward processing/impulsivity, cognitive flexibility/monitoring, inhibition) will be stronger in the FH+ group, compared to the FH- group.

## Methods

### Participants

The current study utilized data from the Human Connectome Project (WU-MINN HCP 1200 Subjects data release), an open-access data initiative which provides comprehensive imaging, behavioral, and genetic data for 1200 healthy subjects aged 22-35. Each research participant was administered the same research protocol and scanned on the same machine. The HCP excludes individuals with documented neuropsychiatric disorders, neurologic disorders, diabetes, high blood pressure, premature birth, and severe symptoms associated with substance use (Van Essen *et al.* 2013). Participants were excluded from present analyses if they did not complete DTI scanning sessions.

### Matched groups

Two groups of participants were selected from the full HCP dataset with complete DTI data, utilizing the case-control matching function in SPSS (v.24). Demographic information for final groups (*N* = 218; 41.3% female) can be found in Supplemental Table 1. Group indicator was defined as self-reported history of chemical dependency in at least one parent. Groups were matched one-to-one on sex, age in years, and diagnostic history of alcohol abuse or dependence (per the Semi-Structured Assessment for the Genetics of Alcoholism). Current alcohol usage (i.e., past 7 days) was defined categorically as “abstinent” (0 drinks), “moderate” (1 – 20 drinks), and “heavy” drinkers (≥ 21 drinks) consistent with previous literature (Makris *et al.* 2008). Education was also defined categorically (11-12 years, 13-15 years, and ≥ 16 years of education). Participants were then matched exactly on current alcohol usage and education categories. Of note, although groups were not matched for income or other substance usage, pre-analyses revealed no significant group differences on income, illicit drug lifetime usage, current tobacco use, current cannabis use, history of cannabis abuse or dependence. The final groups had an average age of 28.90 (*SD* = 3.67), approximately 14 years of education, and 13.8% had a lifetime history of an alcohol use disorder.

**Table 1.** Summary of interaction effects between measures of white matter connectivity (mean and standard deviation in NQA, GFA, and ISO). Significant findings are separated by task, and hemisphere of finding is denoted with L and/or R (X for the corpus callosum). All findings *p < 0.05* after correcting for tract comparisons. DCCS: Dimension Change Card Sort, DD: Delayed Discounting.

|  |  | **DCCS** |  | **Flanker** |  | **DD** |  |
| --- | --- | --- | --- | --- | --- | --- | --- |
|  |  | *M* | *SD* | *M* | *SD* | *M* | *SD* |
| **NQA** |  |  |  |  |  |  |  |
|  | *Corpus Callosum* |  |  |  | X |  |  |
|  | *Cortico-striatal* |  | L | L | L/R |  |  |
|  | *Cortico-thalamic* |  | L | L | L |  |  |
|  | *Cerebellum* |  |  |  |  |  |  |
|  | *Inferior Longitudinal Fasciculus* |  |  | L/R | L |  |  |
|  | *U Fibers* |  |  | L | L |  |  |
| **GFA** |  |  |  |  |  |  |  |
|  | *Corpus Callosum* |  |  |  |  |  |  |
|  | *Cortico-striatal* |  | L/R | L |  |  |  |
|  | *Cortico-thalamic* |  | L/R | L |  |  |  |
|  | *Cerebellum* |  |  |  |  |  | L |
|  | *Inferior Longitudinal Fasciculus* |  |  | R |  |  | R |
|  | *U Fibers* |  | L |  |  |  | R |
| **ISO** |  |  |  |  |  |  |  |
|  | *Corpus Callosum* |  |  | X |  |  | X |
|  | *Cortico-striatal* |  |  | R |  |  |  |
|  | *Cortico-thalamic* |  |  | R | R |  | R |
|  | *Cerebellum* |  |  |  |  |  |  |
|  | *Inferior Longitudinal Fasciculus* |  |  |  |  |  | R |
|  | *U Fibers* |  | R | R | R |  |  |

### Diffusion Tensor Imaging

#### Image acquisition and preprocessing

DTI scans were acquired on a Siemens 3T Skyra system, with a SC72 gradient coil and simultaneous multi-slice echo planar imaging with multiband excitation and multiple receivers at 1.25 mm spatial resolution. DTI scans were pre-processed using the HCP diffusion pipeline, which includes several tools to remove motion and scanner related noise. All images were processed in FSL’s BEDPOSTX (Bayesian Estimation of Diffusion Parameters Obtained using Sampling Techniques, modeling crossing X fibers) to model white matter fiber orientations and crossing fibers for probabilistic tractography. Details on image acquisition, pre-processing, and data quality can be found in previously published literature (Sotiropoulos *et al.* 2013).

#### Reconstruction and group connectometry analysis

Group connectometry analyses (Yeh *et al.* 2016) were performed using DSI-Studio (http://dsi-studio.labsolver.org) to study the effect of family history. Diffusion data was reconstructed in MNI space using q-space diffeomorphic reconstruction (QSDR) (Yeh *et al.* 2010, 2011) to obtain the spin distribution function (SDF) with a sampling length ratio of 1.25. A multiple regression model was used to consider family history for all 218 subjects, controlling for age and sex. A T-score threshold of 3.65 (corresponding to a medium effect size, d ≥ 0.50) was assigned to select local connectomes using a deterministic fiber tracking algorithm (Yeh *et al.* 2016). A false discovery rate (FDR) threshold of 0.05 was used to select tracks showing significant differences between groups, with 1 seed per 0.15 mm3, and two iterations of track trimming. Given previous literature showing the lower bound for major tracts (Hasan *et al.* 2009), subsequent analyses were also completed with track inclusion of at least >40 mm to determine if differences were in major tracts. To estimate the false discovery rate, a total of 5000 randomized permutations were used to obtain the null distribution of tract length. The length is calculated by adding the distance between *consecutive* coordinates along a tract. Network analysis of selected tracks was generated with DSI Studio after permutation analysis. Of note, differences from group connectometry analyses identify continuous segments with significant differences in SDF at peak orientation, regardless of whether those segments make up an entire tract. This is in contrast to traditional fiber tracking, which identifies differences in averages across the entire length of a tract.

### Tract-Behavior Relationships

#### Cognitive Tasks

As part of the behavioral data research protocol, participants completed cognitive tasks included in the NIH Toolbox (http://www.nihtoolbox.org/). Current analyses included data from the following cognitive tasks expected to be related to alcohol and substance use vulnerability: Delayed Discounting, Dimensional Change Card Sort, Flanker Inhibitory Control and Attention Task, and Oral Reading Recognition as a comparison task. These cognitive tasks measure domains of reward processing/impulsivity, cognitive flexibility/monitoring, inhibition, and language/reading decoding. Further details about these cognitive tasks has been provided in previous literature (Weintraub *et al.* 2013; Van Essen *et al.* 2013).

#### Measures of White Matter Microstructure

The following measures of white matter microstructure of selected tracts were extracted for further analysis (Hypothesis 2): normalized quantitative anisotropy (NQA; normalized to the maximum QA in the brain for between-subject comparisons, where QA is the SDF of the resolved fiber orientation minus the background isotropic component), generalized fractional anisotropy (GFA), and isotropic diffusion component (ISO). Tracts were identified using the HCP-842 Atlas available in DSI-Studio for QSDR reconstructed tracts. Extracted values represent the mean and standard deviation along the *entire tract length* of a single subject. Both mean and standard deviation were extracted for analyses because SDF metrics have high variability between- and within-subjects, and variability within structures may show meaningful differences as it does in grey matter brain structures (Wierenga *et al.* 2018). Regarding interpretation, NQA is a density measurement of anisotropy, and thus related to the compactness, or restriction of spins, within fibers along the continuous length of a tract and axonal density. Scaled 0 to 1, values closer to 1 represent more compact fibers. In contrast, while ISO is another density measurement, it characterizes background diffusion, and is thus more related to global diffusion factors such as vasogenic tissue edema (Chiang *et al.* 2014). These were estimated by taking the minimum value of the SDF. Finally, GFA is highly correlated with traditional fractional anisotropy. It describes the preferential directional diffusion mobility, and is a refinement of the traditional approach, but accommodates more complex diffusion information (Glenn *et al.* 2015).

#### Statistical Analyses

Partial correlation matrices, controlling for age and sex, were generated between cognitive tasks and measures of white matter microstructure in tracts that were identified as having *significant group differences*. Correlation matrices for each group (FH+ and FH-) were then compared using a Fisher’s r-to-z-transformation to determine if the relationship between tract characteristics and behavior was moderated by family history of chemical dependency. To further determine whether the relationship between white matter and behavioral measures varied by group, we utilized a general linear model with non-parametric permutation testing (5000 permutations) via FSL’s PALM to account for family structure in the HCP data. Interactions were tested with a two-group regression model with continuous covariate interaction, adjusted for main effects, sex, multiple tract comparisons, and family structure (*p* < .05).

## Results

### Effects of Family History

#### Family History

In regression models controlling for sex and current drinking, there were significant main effects for family history on the local connectome. Using an FDR cut-off, the FH+ group (*N* = 109) had significantly lowered connectivity compared to the FH- group (*N* = 109), in 1092 segments of bilateral cerebellum, bilateral u fibers, bilateral cortico-striatal, left cortico-thalamic, left inferior longitudinal fasciculus and the corpus callosum (FDR < 0.05; See Supplemental Figure 1). Restricting tract selection to major tracts (> 40mm), analyses revealed that the FH+ had significantly lowered connectivity in 97 segments (*M* = 46.69 mm, *SD* = 6.06 mm) of left inferior longitudinal fasciculus, bilateral cortico-striatal pathway, left cortico-thalamic pathway, and corpus callosum (FDR < 0.05; See Figure 1). Fibers in the corpus callosum were localized to interhemispheric, posterior cingulate regions. For these major tracts, network property analyses revealed that these group differences accounted for a 16.1% decrease in overall density, a 13.1% decrease in the clustering coefficient, and 11.6% decrease in small worldedness, and a 13.9% decrease in local efficiency for the FH+ group. It is important to note that the bilateral U fibers, or short association fibers which connect adjacent brain regions, would not be expected to reach sufficient lengths to be characterized as “major tracts”. There is significant variability within these tracts, and while visualization revealed differences in continuous segments between cortical, cerebellar, and striatal regions implicated in reward signaling, the average quantitative anisotropy across the entire length of tracts were nearly identical between groups.

**Figure 1.**
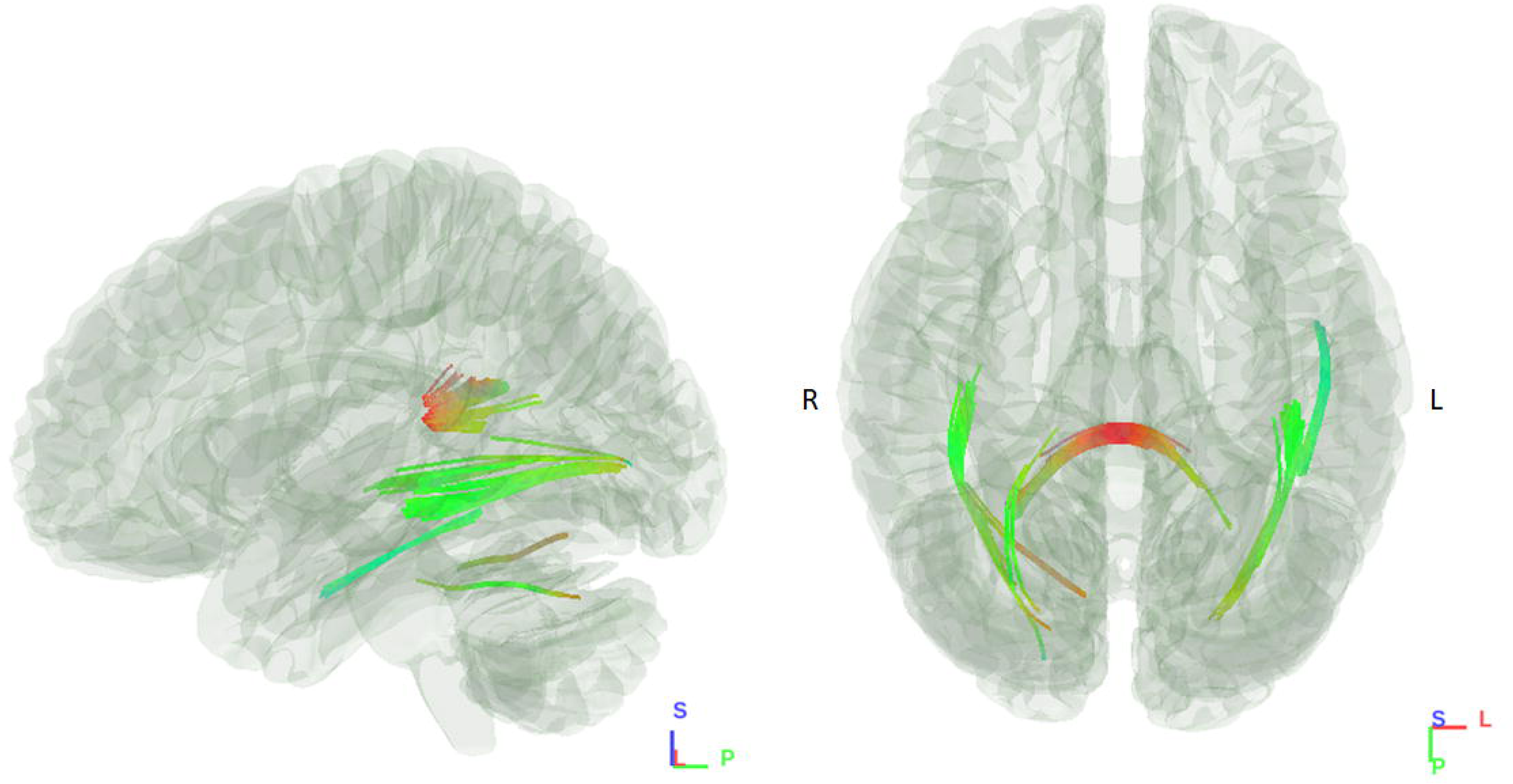
Major tract findings, where the FH+ group had significantly lowered connectivity in segments of at least 40mm or greater, compared to the FH- group.

#### Sex Effects

Post-hoc analyses were performed to determine if the two sexes were differentially affected by family history. There were no interactions between family history and sex in group connectometry analyses, using either FDR or tract length thresholds. For all subsequent analyses of tract-behavior relationships, sex was controlled for in addition to age as a covariate of non-interest.

#### Tract-Behavior Relationships

Between groups, there were no significant differences in performance on cognitive measures, or on measures of NQA, GFA, or ISO within specific identified tracts (See Supplemental Table 2). In contrast to local connectome analyses, NQA, GFA, and ISO values represent the average value along the *full* length of the tract. However, for some tasks, tracts, and measures there was a significant *interaction*, where the association between brain and behavior differed between groups.

Generally, when the relationship between brain and behavior significantly differed between groups, the association was stronger in FH+ compared to FH-, although there was some variability by task and measure. For the reward processing task (i.e., Delayed Discounting), performance was positively associated with GFA and ISO measures, negatively with NQA measures, and generally stronger in the FH+ group. For the inhibition task (i.e., Flanker), measures of GFA and NQA were weakly, positively associated with performance in the FH- group, and weakly, negatively associated with performance in the FH+ group. However, the reverse was true for the measures of ISO. A similar pattern emerged for monitoring task (i.e., Card Sort), with the exception of mean GFA, which was weakly negatively associated with performance in the FH- group. As expected, the relationship between tracts and our comparison task (i.e., Reading) did not significantly vary between groups on any measure after permutations and correction for multiple comparisons. Reading was positively associated with measures of GFA and ISO and negatively with measures of NQA. While the differences in interaction effects size between the comparison task and tasks of interest were largely non-significant (i.e., overlapping confidence intervals), the consistency in results for the comparison task is notable.

Results for tract specific findings are described separately for NQA and GFA measures and the three behavioral tasks of interest (i.e., reward processing, monitoring, inhibition). Tract specific findings for ISO are presented in Supplement 1. A summary of the interaction findings is shown in Table 1 and representative scatter plots are shown in Figure 2 to demonstrate the patterns of effects. Partial correlation matrices separated by group for mean and standard deviation of NQA, GFA, and ISO along the full length of white matter microstructure are available in Supplemental Tables 3-8.

**Figure 2.**
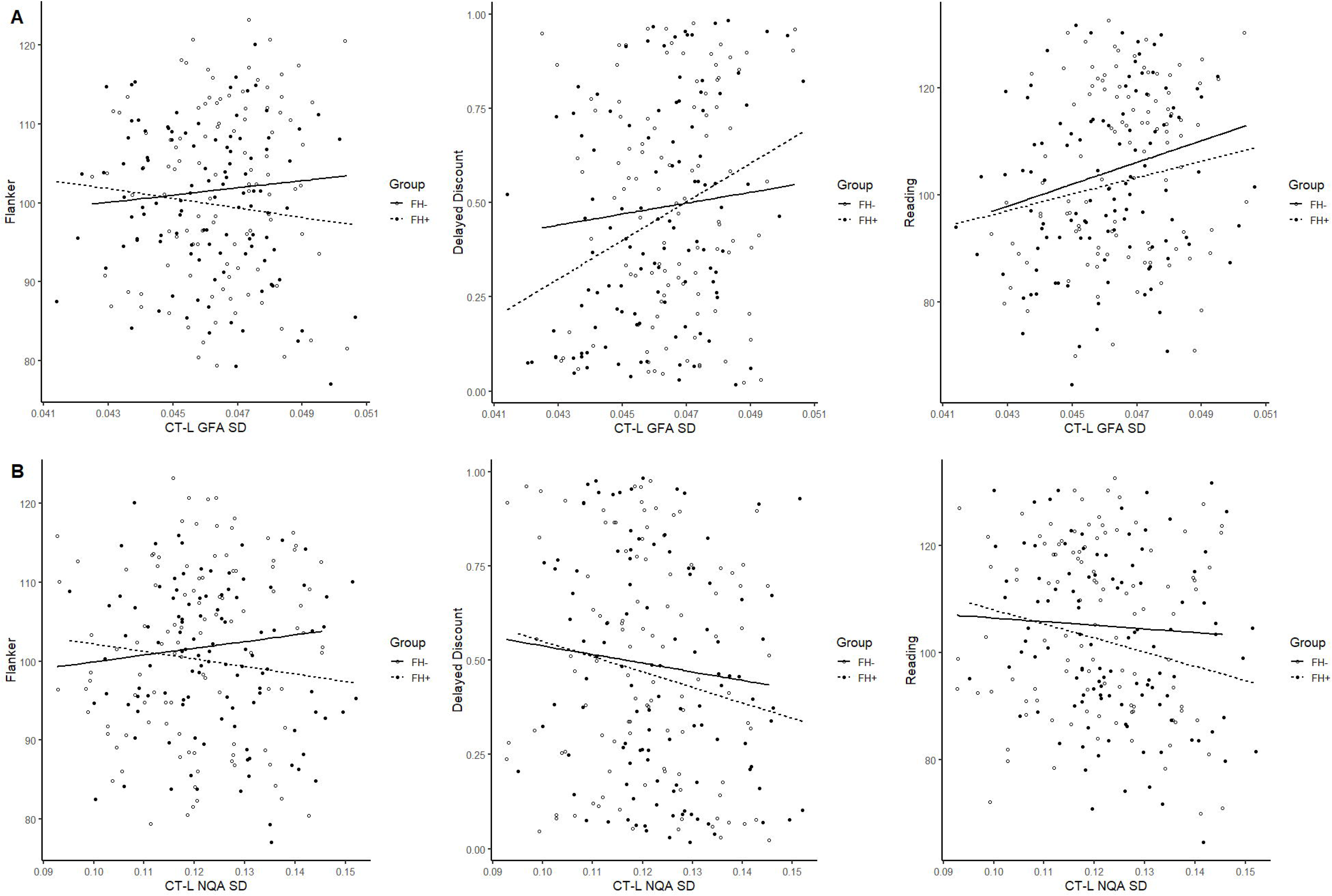
Representative scatter plots of the brain, behavior, group interaction. The following tasks are shown on the y-axis: Flanker (Standard Score), Delayed Discounting (Area under the curve), and Reading (Standard Score). The cortico-thalamic pathway is represented on the x-axis for measures of within tract GFA (A) and NQA (B) *variability*. Dimensional Card Sort is not shown, as the patterns of findings were similar to the Flanker.

#### NQA

Mean NQA was significantly negatively associated with reward processing in bilateral cortico-striatal pathways, left cortico-thalamic pathways, left u-fibers, and the corpus callosum, and with reading in the right cortico-thalamic and cortico-striatal pathways, for FH+ individuals only. However, there was no significant interaction, suggesting that the brain-behavior relationship did not differ between groups. The interaction analyses did reveal that the relationship between inhibition and bilateral inferior longitudinal fasciculi, left u-fibers, left cortico-thalamic pathway, and the left cortico-striatal pathway differed between groups, with weak negative correlations in the FH+ group and weak positive correlations in the FH- group.

NQA variability was also significantly negatively associated with reward processing and reading in the corpus callosum, left cortico-thalamic, and –striatal pathways for FH+. Again, these findings were not found to be significant within the interaction analyses. For both the inhibition and cognitive flexibility/monitoring tasks, the brain-behavior relationship significantly varied between groups, with similar patterns of directional flipping as in the mean NQA findings. NQA variability associations with inhibition differed in the left inferior longitudinal fasciculus, u-fibers, cortico-thalamic pathways, corpus callosum, and bilateral cortico-striatal pathways while the association with monitoring only differed in the left cortico-striatal, and cortico– thalamic pathways.

#### GFA

Generally, the average GFA of identified tracts was not associated with performance on reward processing tasks in either group. In contrast, variability of GFA within identified tracts was positively associated with tasks of reward processing for the FH+ group only. However, after testing the interaction in PALM only GFA variability in left cerebellum, right inferior longitudinal fasciculus, and right u-fibers was more significantly related to reward processing in the FH+ group.

For cognitive flexibility/monitoring, performance was not significantly related to GFA averages or variability in almost every tract. However, when examining the interaction, the relationship between brain and behavior significantly differed between groups for variability in bilateral cortico-thalamic pathway, cortico-striatal pathways, and left u-fibers. Association with average GFA differed between groups for the inhibition task only, in the left cortico-striatal pathway, left cortico-thalamic pathways, and right inferior longitudinal fasciculus.

## Discussion

The principal findings of this study are that (1) family history of drug and alcohol dependence was associated with decreased white matter connectivity along ventral and subcortical tracts in adulthood, including (2) major tracts in the left cortico-thalamic, bilateral cortico-striatal, left inferior longitudinal fasciculus and corpus callosum. Additionally, (3) while across the entire length of the tracts, average density based measurements were nearly identical, (4) the relationship between white matter microstructure and clinically relevant behavior was significantly different between groups. These results are impactful for a number of reasons.

This is the first study to leverage the advantages of density based white matter measurements to examine structural vulnerabilities conferred by a family history of chemical dependency. All significant identified segments of major tracts were found within the ventral and subcortical pathways associated with reward and inhibitory processing. Specifically, these tracts connect regions associated with cue reactivity in frequent drinkers (Fryer *et al.* 2013) and projections may be associated with saccade reward signals and saliency given the involvement of the posterior cingulate cortex in those functions (McCoy *et al.* 2003; Vossel *et al.* 2014). These results are also consistent with previous dMRI studies in individuals with severe alcohol misuse symptomatology (Seitz *et al.* 2017; Sawyer *et al.* 2018). The major tract identification finding of the interhemispheric connections are consistent with *functional findings* in adolescents (Herting *et al.* 2011), of dysregulation in contra-lateral cortico-subcortical projections. While the findings in that study identified corresponding decreases in fractional anisotropy in the superior longitudinal fasciculus and internal capsule, they did not identify interhemispheric tracts that may impact contralateral connectivity. This may have resulted from the increased proportion of crossing fibers in the corpus callosum. The novel identification of u-fibers as an area of vulnerability for this population may result from their relative “sparing” or delayed involvement in pathological processes, due to their resistance to myelin metabolic effects. U-fibers are often impacted when oligodendroglia cell bodies are directly affected, as opposed to myelin metabolic turnover (Welker & Patton 2012; Riley *et al.* 2018), and these cell body changes may not have yet impacted the integrity of expressed myelin. A density based measure, which more accurately represents the axonal integrity, is therefore more sensitive to the effects of u-fibers particularly in subclinical populations.

Our findings are in young adults, where we would expect white matter to be more fully developed relative to previous studies of adolescents. This supports the conclusion that vulnerabilities conferred by a family history of chemical dependency persist into adulthood, and do not simply represent a slowed trajectory of white matter development. While it is possible that slowed development contributed to riskier behaviors in adolescence, and subsequent damage to white matter persisting into adulthood, our stringent matching criteria between groups make this explanation less likely. The age of our sample may also contribute to the increased interaction findings in the left hemisphere for the cognitive flexibility/monitoring task, which is consistent with theories of compensatory mechanisms and lowered bilateral recruitment needs in early adulthood (Reuter-Lorenz & Cappell 2008). Through early and middle adulthood, these vulnerabilities may compound with increasing environmental stress and recruitment needs, and examinations of familial risk should consider the role of these biological differences throughout the entire lifespan developmental trajectory. Similarly, while we did not find interactive sex differences in this sample, a lifespan approach may be impacted by hormonal differences in the latter half of the lifespan and interact with drinking behavior (Seitz *et al.* 2017; Sawyer *et al.* 2018). However, to fully understand these lifespan trajectories, longitudinal research is needed in the population which extend from adolescence into early and middle adulthood.

Our findings suggest that the relationship between brain and behavior differed significantly between groups. While many of the relationships between white matter and inhibitory or monitoring processes were non-significant, there were a number of significant interactive effects across NQA, GFA, and ISO. In particular, the number of interaction effects found for the monitoring and inhibition tasks imply that systems involved in attending to reinforcement and error monitoring may be especially vulnerable. The patterns of findings were also similar for these two tasks regarding strength and direction, perhaps due to the involvement of error monitoring across both. Given the relative health of this population, and the lack of group differences on behavioral tasks or tract quality, we believe these interactive effects reveal underlying vulnerabilities that could emerge over time. The consistency of the comparison task across groups support our conclusion that these vulnerabilities differentially affect cognitive skills in reward and inhibitory processes. Family history of chemical dependency does not confer global abnormalities, but rather specific dysregulation that contribute to the future risk of alcohol and substance misuse. This makes it more likely that these differences result from the genetic contributions of family history, and not associated social factors that affect more global measures, and which may be more prevalent among parents with alcohol and substance use problems.

The discrepancy in correlational direction in the NQA and GFA findings is notable. As expected, increased mean GFA was generally positively associated with reward processing performance across both groups, which is consistent with findings in traditional diffusion measures (Samanez-Larkin *et al.* 2012). This relationship was reversed for mean NQA in the FH+ group, suggesting as fiber density approaches maximum values, performance on tasks of reward processing becomes worse. This relationship may reflect a dysregulation in *axonal body* pruning, which is supported by findings which show that increased variability in NQA (i.e., uneven or unreliable pruning) is also associated with worse performance. Comparatively, variability in GFA was associated with better performance and may be more reflective of the *myelin* integrity that is constantly remodeling within healthy adults in ways that axon bodies cannot (Young *et al.* 2013). However, because larger ISO values (or minimum SDF) were associated with better performance in the FH+ group, the impact of pruning on performance may follow a Yerkes–Dodson distribution. As with findings on risk taking behaviors and increased fractional anisotropy (Berns *et al.* 2009; Squeglia *et al.* 2014), these differences may also be associated with maladaptive behaviors outside the scope of neuropsychological testing measures. Notably, density based measures also displayed lateralized trends within analyses for behavioral relevance, where NQA association with behavior largely differed between groups in the left hemispheric major tracts, while ISO associations differed largely in the right hemisphere. This suggests there may also be differential effects of family history on hemispheres in early adulthood. More research is needed to further explore these dissociations.

There are several important limitations in this study which present the potential for future investigations. Our goal was to examine the effects on white matter when at a more developmentally advanced stage compared to earlier research, but we recognize that there is individual variation for neural developmental trajectories. Further, our focus on early adulthood has limited the impact of our findings from a lifespan perspective. Future investigations would benefit from examining the impact of family history on density-based measurements across the full lifespan to better characterize the trajectory and risk associated with these vulnerabilities. These findings largely describe biological characteristics that have not yet resulted in observable behavioral outcomes, and are therefore limited in their clinical relevance at the present time. However, we believe that future investigations of the interaction between family history and substance use may reveal more clinically useful information regarding critical time points of intervention.

In summary, to our knowledge this is the first study examining the effect of family history of alcohol and substance use utilizing a group connectometry analysis. Density based measurements were more sensitive to axonal body differences in white matter, particularly in areas where fibers cross. Our findings suggest that these vulnerabilities may exist in early adulthood, are likely the end result of maturational differences identified in previous studies of adolescent samples, and are specific to reward and inhibitory processes. Further investigation utilizing density based measures will increase our understanding of the mechanisms by which family history affects neural characteristics will help to identify treatment targets and critical periods of intervention.

## Acknowledgments

The authors would like to acknowledge the following people and organizations for their contributions:

Data were provided [in part] by the Human Connectome Project, WU-Minn Consortium (Principal Investigators: David Van Essen and Kamil Ugurbil; 1U54MH091657) funded by the 16 NIH Institutes and Centers that support the NIH Blueprint for Neuroscience Research; and by the McDonnell Center for Systems Neuroscience at Washington University.

The Suffolk University Psychology Department for their support of doctoral students and David Gansler’s Lab.

---

**Supplemental Figure 2**. All tract findings (FDR < 0.05) for decreased connectivity in the FH+ group compared to FH- group.

