## Supplementary figures and images for "White Matter Connectometry Among Individuals with Self-Reported Family History of Drug and Alcohol Use Disorders"

### Supplemental Figure 1

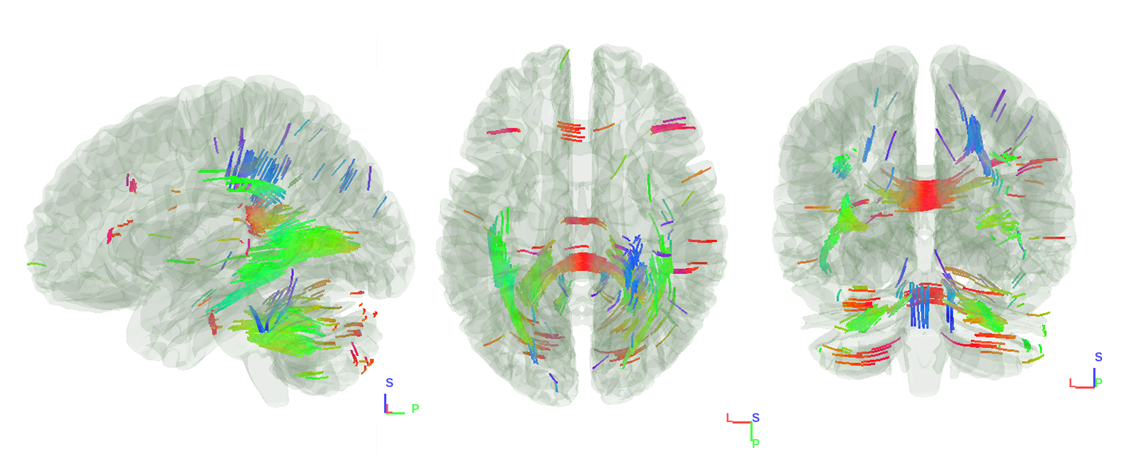
