## Supplement for "White Matter Connectometry Among Individuals with Self-Reported Family History of Drug and Alcohol Use Disorders"

**Supplement 1. Specific Tract Findings for ISO**

**ISO.** For the FH+ group, while average ISO was associated with reward processing in the corpus callosum, bilateral u-fibers, and right cortico-thalamic pathways in the FH+ group, interaction analyses revealed that the relationship between brain and behavior was not significantly stronger in any trast, compared to the FH- group. While mean ISO is largely not significantly associated with the inhibition task, the relationship did differ significantly between groups in the right cortico-thalamic, cortico-striatal, u-fibers, and corpus callosum. Specifically, for the FH- group increased ISO was associated with decreased performance, while the opposite was true for the FH+ group.

Results from the variability with tracts were similar, although there was a lateralization of findings. ISO variability was strongly positively associated with reward processing, differing by group in the right cortico-thalamic pathways, inferior longitudinal fasciculus, and corpus callosum. Again, there were significant interactions in the right cortico-thalamic pathway and u-fibers for the inhibition task, and for right u-fibers in the monitoring task, even though these relationships were largely insignificant as partial correlations.

*Supplemental Table 1*. Demographic variables separated by group. Note that *Income* was coded as follows: <$10,000 = 1,10K-19,999 = 2, 20K-29,999 = 3,30K-39,999 = 4, 40K-49,999 = 5, 50K-74,999 = 6, 75K-99,999 = 7, >=100,000 = 8.

|  | FH + (*N* = 109) | | FH - (*N* = 109) | |
| --- | --- | --- | --- | --- |
|  | *M* | *SD* | *M* | *SD* |
| Age | 28.90 | 3.67 | 28.90 | 3.67 |
| Education (Years) | 14.17 | 1.98 | 14.27 | 1.98 |
| Income | 4.50 | 2.12 | 4.75 | 2.19 |
| ASR Anxiety (T-Score) | 54.37 | 6.23 | 53.18 | 4.72 |
| ASR Depression (T-Score) | 54.50 | 6.55 | 53.63 | 5.90 |
| Total Drinks (Past 7 Days) | 4.18 | 5.98 | 3.80 | 5.91 |
|  | % | | % | |
| Gender (% Female) | 41.3 | | 41.3 | |
| Race |  | | | |
| *Asian/Nat. Hawaiian/Pacific Islander* | 2.8 | | 3.7 | |
| *Native American/Alaskan* | 0.0 | | 0.9 | |
| *Black* | 36.7 | | 15.6 | |
| *White* | 56.9 | | 76.1 | |
| *Other* | 3.7 | | 3.7 | |
| Ethnicity |  | | | |
| *Hispanic/Latino* | 5.5 | | 6.4 | |
| Tobacco Smoker (Current/Lifetime) | 27.5/58.7 | | 22.0/45.9 | |
| Cannabis Smoker (Lifetime) | 62.4 | | 50.5 | |
| Alcohol Use Disorder (Lifetime) | 13.8 | | 13.8 | |
| Cannabis Use Disorder (Lifetime) | 10.1 | | 9.2 | |

*Supplemental* *Table 2*. Cognitive, NQA, GFA, and ISO values for the *full length of tract* separated by group. All bilateral tracts are reported L/R. GFA and NQA are scaled 0 to 1.

|  | FH + (*N* = 109) | | FH - (*N* = 109) | |
| --- | --- | --- | --- | --- |
|  | *M* | *SD* | *M* | *SD* |
| Cognitive Variables |  |  |  |  |
| *Dimensional Card Sort* | 100.39 | 9.91 | 101.88 | 11.01 |
| *Flanker* | 99.95 | 9.30 | 101.69 | 11.48 |
| *Delayed Discounting* | 0.21 | 0.18 | 0.26 | 0.21 |
| NQA |  |  |  |  |
| *Corpus Callosum* | 0.23 | 0.13 | 0.22 | 0.13 |
| *Corticostriatal* | 0.24/0.25 | 0.12/0.11 | 0.23/0.22 | 0.12/0.11 |
| *Corticothalamic* | 0.24/0.23 | 0.12/0.11 | 0.23/0.23 | 0.12/0.11 |
| *Cerebellum* | 0.15/0.14 | 0.05/0.05 | 0.15/0.14 | 0.05/0.05 |
| *Inferior Longitudinal Fasciculus* | 0.24/0.24 | 0.14/0.14 | 0.24/0.24 | 0.14/0.14 |
| *U Fibers* | 0.20/0.18 | 0.12/0.11 | 0.19/0.17 | 0.12/0.11 |
| GFA |  |  |  |  |
| *Corpus Callosum* | 0.10 | 0.05 | 0.10 | 0.05 |
| *Corticostriatal* | 0.10/0.10 | 0.05/0.04 | 0.10/0.10 | 0.05/0.05 |
| *Corticothalamic* | 0.10/0.10 | 0.05/0.05 | 0.10/0.10 | 0.05/0.05 |
| *Cerebellum* | 0.06/0.06 | 0.02/0.02 | 0.06/0.06 | 0.02/0.02 |
| *Inferior Longitudinal Fasciculus* | 0.11/0.10 | 0.05/0.05 | 0.11/0.10 | 0.05/0.05 |
| *U Fibers* | 0.08/0.08 | 0.05/0.05 | 0.08/0.08 | 0.05/0.05 |
| ISO |  |  |  |  |
| *Corpus Callosum* | 1.78 | 0.41 | 1.80 | 0.42 |
| *Corticostriatal* | 1.89/1.72 | 0.43/0.32 | 1.88/1.73 | 0.42/0.32 |
| *Corticothalamic* | 1.84/1.72 | 0.41/0.33 | 1.73/1.83 | 0.34/0.30 |
| *Cerebellum* | 2.19/2.13 | 0.40/0.41 | 2.23/2.17 | 0.41/0.43 |
| *Inferior Longitudinal Fasciculus* | 1.69/1.81 | 0.40/0.47 | 1.72/1.86 | 0.42/0.50 |
| *U Fibers* | 1.83/1.82 | 0.30/0.32 | 1.83/1.83 | 0.30/0.33 |

*Supplemental Table 3.* Partial Correlations controlling for Age and Gender, between cognitive tasks and Mean GFA. Partial correlations were generated using 500 bootstrapped samples, **p ≤ .05, **p ≤ .01*. *Italicized pairs* have non-overlapping confidence intervals, generated by Fisher’s z transformation, while **bolded pairs** have non-overlapping confidence intervals generated by bootstrapped samples.

|  | Task | CS-L | CS-R | CT-L | CT-R | ILF-L | ILF-R | CM-L | CM-R | U-L | UF-R | CC |
| --- | --- | --- | --- | --- | --- | --- | --- | --- | --- | --- | --- | --- |
| Family History  Negative | Delayed Discount | 0.13 | 0.06 | 0.10 | *0.06* | *0.09* | *-0.01* | *-0.05* | *-0.08* | 0.15 | *0.00* | 0.07 |
|  | Card Sort | -0.09 | -0.06 | -0.03 | -0.06 | 0.09 | 0.02 | -0.03 | -0.01 | *0.03* | -0.06 | -0.08 |
|  | Flanker | *0.04* | 0.03 | *0.06* | 0.02 | 0.11 | -0.09 | 0.02 | 0.09 | *0.10* | 0.03 | 0.02 |
|  | Reading | 0.08 | 0.11 | 0.12 | 0.15 | 0.18 | .20* | *-0.01* | *-0.14* | 0.17 | *0.06* | 0.12 |
| Family History  Positive | Delayed Discount | 0.13 | 0.17 | 0.18 | *0.20** | *.32*** | *.26*** | *0.11* | *0.07* | 0.18 | *0.17* | 0.19 |
|  | Card Sort | -0.17 | -0.16 | -0.16 | -0.14 | -0.05 | -0.08 | -0.02 | 0.00 | *-0.13* | -0.07 | -0.12 |
|  | Flanker | *-0.17* | -0.08 | *-0.14* | -0.07 | 0.03 | 0.03 | 0.00 | -0.03 | *-0.09* | 0.00 | -0.05 |
|  | Reading | 0.16 | 0.16 | 0.21* | 0.18 | .32** | .29** | *0.14* | *0.14* | 0.26** | *0.25*** | 0.22* |

|  | Task | CS-L | CS-R | CT-L | CT-R | ILF-L | ILF-R | CM-L | CM-R | U-L | UF-R | CC |
| --- | --- | --- | --- | --- | --- | --- | --- | --- | --- | --- | --- | --- |
| Family History  Negative | Delayed Discount | *0.08* | *0.05* | *0.12* | *0.11* | *0.08* | *0.12* | *-0.07* | *0.07* | *0.16* | *0.06* | *0.07* |
|  | Card Sort | *0.10* | *0.17* | *0.17* | *0.17* | 0.12 | 0.08 | *-0.10* | *-0.12* | *0.20** | *0.14* | *0.08* |
|  | Flanker | *0.07* | *0.09* | *0.09* | 0.09 | -0.01 | 0.1 | -0.06 | -0.01 | *0.17* | 0.07 | 0.05 |
|  | Reading | 0.26** | 0.22* | 0.25** | 0.22* | .31** | .34** | *0.00* | *-0.06* | 0.20* | 0.23* | 0.20* |
| Family History  Positive | Delayed Discount | *0.22** | *0.29*** | *0.27*** | *0.33*** | *.31*** | *.20** | *0.26*** | *0.24** | *0.30*** | *0.31*** | *0.23** |
|  | Card Sort | *-0.13* | *-0.12* | *-0.08* | *-0.11* | -0.04 | 0.01 | *0.06* | *0.06* | *-0.07* | *-0.08* | *-0.06* |
|  | Flanker | *-0.13* | *-0.06* | *-0.08* | -0.03 | 0.08 | 0.12 | 0.02 | 0.06 | *-0.04* | 0.09 | 0.02 |
|  | Reading | 0.14 | 0.15 | 0.19 | 0.18 | .32** | .21* | *0.24** | *0.28*** | 0.23* | 0.25** | 0.19* |

*Supplemental Table 4.* Partial Correlations controlling for Age and Gender, between cognitive tasks and SD GFA. Partial correlations were generated using 500 bootstrapped samples, **p ≤ .05, **p ≤ .01*. *Italicized pairs* have non-overlapping confidence intervals, generated by Fisher’s z transformation, while **bolded pairs** have non-overlapping confidence intervals generated by bootstrapped samples.

|  | Task | CS-L | CS-R | CT-L | CT-R | ILF-L | ILF-R | CM-L | CM-R | U-L | UF-R | CC |
| --- | --- | --- | --- | --- | --- | --- | --- | --- | --- | --- | --- | --- |
| Family History  Negative | Delayed Discount | *-0.02* | *-0.03* | *-0.01* | *-0.01* | 0.08 | 0.03 | -0.01 | -0.02 | *-0.01* | -0.03 | *0.00* |
|  | Card Sort | 0.04 | 0.04 | 0.05 | 0.03 | 0.13 | 0.04 | 0.06 | 0.08 | 0.05 | 0.01 | 0.03 |
|  | Flanker | *0.09* | 0.06 | *0.10* | 0.05 | 0.12 | 0.01 | 0.07 | 0.07 | *0.11* | 0.05 | 0.07 |
|  | Reading | -0.06 | *-0.07* | -0.06 | *-0.06* | 0.01 | 0.07 | -0.04 | -0.05 | -0.06 | -0.09 | -0.06 |
| Family History  Positive | Delayed Discount | *-0.25** | *-0.20** | *-0.24** | *-0.18* | -0.06 | -0.02 | -0.09 | -0.12 | *-0.21** | -0.16 | *-0.20** |
|  | Card Sort | -0.05 | -0.05 | -0.02 | -0.02 | 0.07 | 0.05 | 0.13 | 0.07 | 0.01 | 0.02 | -0.02 |
|  | Flanker | *-0.08* | -0.04 | *-0.07* | -0.03 | 0.03 | 0.05 | 0.06 | 0.01 | *-0.04* | 0.01 | -0.04 |
|  | Reading | -0.19 | *-0.21** | -0.19 | *-0.20** | -0.07 | -0.07 | -0.12 | -0.17 | -0.17 | -0.16 | -0.19 |

*Supplemental Table 5.* Partial Correlations controlling for Age and Gender, between cognitive tasks and Mean NQA. Partial correlations were generated using 500 bootstrapped samples, **p ≤ .05, **p ≤ .01*. *Italicized pairs* have non-overlapping confidence intervals, generated by Fisher’s z transformation, while **bolded pairs** have non-overlapping confidence intervals generated by bootstrapped samples.

|  | Task | CS-L | CS-R | CT-L | CT-R | ILF-L | ILF-R | CM-L | CM-R | U-L | UF-R | CC |
| --- | --- | --- | --- | --- | --- | --- | --- | --- | --- | --- | --- | --- |
| Family History  Negative | Delayed Discount | *-0.08* | -0.08 | -0.06 | -0.03 | 0.08 | 0.07 | 0.00 | 0.02 | -0.04 | -0.04 | *-0.05* |
|  | Card Sort | *0.08* | *0.08* | *0.11* | *0.07* | 0.12 | 0.06 | 0.09 | 0.10 | *0.10* | 0.05 | *0.07* |
|  | Flanker | *0.13* | *0.09* | *0.14* | *0.09* | 0.06 | 0.07 | 0.07 | 0.06 | *0.15* | 0.07 | *0.12* |
|  | Reading | *-0.03* | *-0.04* | *-0.03* | -0.03 | 0.11 | 0.15 | 0.01 | 0.01 | -0.05 | -0.03 | *-0.05* |
| Family History  Positive | Delayed Discount | *-0.23** | -0.13 | -0.22* | -0.05 | 0.06 | 0.06 | 0.06 | 0.02 | -0.17 | -0.08 | *-0.21** |
|  | Card Sort | *-0.08* | *-0.08* | *-0.07* | *-0.07* | 0.06 | 0.04 | 0.14 | 0.04 | *-0.05* | -0.05 | *-0.07* |
|  | Flanker | *-0.11* | *-0.08* | *-0.11* | *-0.07* | 0.08 | 0.07 | 0.03 | 0.03 | *-0.08* | 0.01 | *-0.06* |
|  | Reading | *-0.21** | *-0.19** | *-0.20** | -0.16 | 0.05 | 0.06 | 0.02 | -0.01 | -0.18 | -0.13 | *-0.21** |

*Supplemental Table 6.* Partial Correlations controlling for Age and Gender, between cognitive tasks and SD NQA. Partial correlations were generated using 500 bootstrapped samples, **p ≤ .05, **p ≤ .01*. *Italicized pairs* have non-overlapping confidence intervals, generated by Fisher’s z transformation, while **bolded pairs** have non-overlapping confidence intervals generated by bootstrapped samples.

*Supplemental Table 7.* Partial Correlations controlling for Age and Gender, between cognitive tasks and Mean ISO. Partial correlations were generated using 500 bootstrapped samples, **p ≤ .05, **p ≤ .01*. *Italicized pairs* have non-overlapping confidence intervals, generated by Fisher’s z transformation, while **bolded pairs** have non-overlapping confidence intervals generated by bootstrapped samples.

|  | Task | CS-L | CS-R | CT-L | CT-R | ILF-L | ILF-R | CM-L | CM-R | U-L | UF-R | CC |
| --- | --- | --- | --- | --- | --- | --- | --- | --- | --- | --- | --- | --- |
| Family History  Negative | Delayed Discount | -0.13 | *-0.14* | *-0.12* | *-0.09* | *0.03* | *0.03* | *-0.01* | *-0.02* | *-0.12* | *-0.08* | *-0.06* |
|  | Card Sort | 0.07 | 0.02 | 0.06 | -0.02 | 0.09 | -0.01 | 0.01 | 0.02 | *0.00* | *-0.04* | 0.01 |
|  | Flanker | -0.07 | *-0.18* | *-0.10* | *-0.20** | -0.08 | -0.12 | *-0.09* | *-0.11* | *-0.08* | *-0.18* | *-0.16* |
|  | Reading | 0.07 | 0.06 | 0.09 | 0.06 | 0.16 | 0.14 | 0.11 | 0.13 | *0.04* | 0.04 | 0.11 |
| Family History  Positive | Delayed Discount | -0.01 | *0.16* | *0.07* | *0.25*** | *.24** | *.24** | *0.19* | *0.17* | *0.16* | *0.26*** | *0.24** |
|  | Card Sort | -0.01 | 0.00 | 0.04 | 0.05 | 0.08 | 0.08 | 0.13 | 0.07 | *0.15* | *0.12* | 0.08 |
|  | Flanker | 0.04 | *0.10* | *0.06* | *0.10* | 0.08 | 0.08 | *0.10* | *0.07* | *0.14* | *0.14* | *0.10* |
|  | Reading | 0.11 | 0.14 | 0.18 | 0.18 | .20* | 0.15 | 0.13 | 0.09 | *0.22** | 0.18 | 0.21* |

*Supplemental Table 8.* Partial Correlations controlling for Age and Gender, between cognitive tasks and SD ISO. Partial correlations were generated using 500 bootstrapped samples, **p ≤ .05, **p ≤ .01*. *Italicized pairs* have non-overlapping confidence intervals, generated by Fisher’s z transformation, while **bolded pairs** have non-overlapping confidence intervals generated by bootstrapped samples.

|  | Task | CS-L | CS-R | CT-L | CT-R | ILF-L | ILF-R | CM-L | CM-R | U-L | UF-R | CC |
| --- | --- | --- | --- | --- | --- | --- | --- | --- | --- | --- | --- | --- |
| Family History  Negative | Delayed Discount | -0.08 | *-0.11* | -0.10 | ***-0.08*** | *0.06* | *0.09* | *-0.02* | *-0.03* | *-0.18* | ***-0.06*** | *-0.11* |
|  | Card Sort | 0.12 | 0.04 | 0.13 | *-0.05* | 0.07 | -0.01 | 0.06 | 0.12 | 0.10 | *-0.06* | 0.05 |
|  | Flanker | 0.00 | *-0.14* | 0.01 | *-0.17* | -0.07 | -0.1 | *-0.06* | *-0.08* | 0.00 | *-0.14* | *-0.10* |
|  | Reading | 0.06 | *0.03* | 0.05 | *0.02* | .23* | .19* | 0.06 | 0.13 | *-0.02* | *0.02* | 0.11 |
| Family History  Positive | Delayed Discount | -0.09 | *0.16* | -0.07 | ***0.30***** | *.26*** | *.33*** | *0.16* | *0.12* | *0.07* | ***0.33***** | *0.21** |
|  | Card Sort | 0.01 | 0.09 | 0.03 | *0.14* | 0.09 | 0.12 | 0.08 | 0.00 | 0.16 | *0.19** | 0.16 |
|  | Flanker | 0.02 | *0.09* | 0.03 | *0.10* | 0.1 | 0.05 | *0.09* | *0.09* | 0.11 | *0.14* | *0.12* |
|  | Reading | 0.05 | *0.17* | 0.05 | *0.20** | .20* | .21* | 0.08 | 0.10 | *0.13* | *0.20** | 0.22* |
